# Equitable Thresholding and Clustering (ETAC): A novel method for FMRI clustering in AFNI

**DOI:** 10.1101/295931

**Authors:** Robert W. Cox

## Abstract

This paper describes a hybrid method to threshold FMRI group statistical maps derived from voxelwise second-level statistical analyses. The proposed “Equitable Thresholding and Clustering” (ETAC) approach seeks to reduce the dependence of clustering results on arbitrary parameter values by using multiple sub-tests, each equivalent to a standard FMRI clustering analysis, to make decisions about which groups of voxels are potentially significant. The union of these sub-test results decides which voxels are accepted. The approach adjusts the cluster-thresholding parameter of each sub-test in an equitable way, so that the individual false positive rates (FPRs) are balanced across sub-tests to achieve a desired final FPR (e.g., 5%). ETAC utilizes resampling methods to estimate the FPR, and thus does not rely on parametric assumptions about the spatial correlation of FMRI noise. The approach was validated with pseudo-task timings in resting state brain data. Additionally, a task FMRI data collection was used to compare ETAC’s true positive detection power vs. a standard cluster detection method, demonstrating that ETAC is able to detect true results and control false positives while reducing reliance on arbitrary analysis parameters.

## INTRODUCTION

One of the challenges in analyzing an FMRI experiment is an appropriate correction for the large number of multiple comparisons, resulting from separate statistical tests at each spatial location (voxel). One strategy for multiple comparisons correction is to leverage the observation that plausible results are likely to span multiple contiguous voxel locations, leading to the rejection of statistical findings that are spatially too small or too insignificant. Leveraging contiguity is commonly carried out using *dual thresholding* of statistical parametric maps, performed in two successive steps. First, examine each voxel and reject any whose test statistic likelihood (p-value) is larger than some user-selected *p*-threshold (referred to herein as “S” or statistical threshold). Second, among the surviving voxels, accept only those that form neighborhoods with other surviving voxels in a cluster of some threshold size or larger (referred to as “C” or clustering threshold).

Both the “S” and “C” steps involve the selection of arbitrary parameters for voxel-wise significance and cluster threshold. The method presented here, called “equitable thresholding and clustering” (ETAC), was developed to reduce the influence of the selection of these arbitrary analysis parameters. ETAC combines the results from using multiple sets of parameters to give experimental results that are not strongly dependent on arbitrary parameter choices. This combination is relevant to concerns about rigor and reproducibility, because it reduces the temptation for researchers to experiment with many different parameter combinations to find those that produce the desired results (even though they may not accurately portray the results that would be found with most parameter values). This effect has been referred to as “the garden of forking paths” problem (Gelman and Loken, 2013).

The complex interdependencies between the “S” and “C” parameters make this problem more complicated than it might appear at first. Typically, S is (somewhat arbitrarily) chosen by the researcher, and then C is estimated for S so that the final map of surviving voxels meets desired statistical stringency (e.g., *p*<0.05). The appropriate C for a given S depends on the smoothness of the data, which depends on arbitrary choices about the size of spatial blurring during image preprocessing. A common sense approach would be to base the choice of both parameters on the size of the region of interest, so that researchers interested in smaller brain areas would apply less smoothing and a more stringent S, so that a small cluster would be able to survive. However, no uniform values for S and C exist in the literature, and any choice within a reasonable range is accepted without further justification. Problems with this approach were brought to the attention of the FMRI community in 2016, with a paper (Eklund et al., 2016) demonstrating that some methods for determining C given S showed markedly higher-than-nominal FPR rates, and this result was dependent on the voxelwise thresholds (S) chosen (Cox et al. 2017a,b).

Finding threshold C from a given S does not have an exact closed-form solution, and there are multiple computational approaches available to find an appropriate C. While some formula-based approaches have been deployed, such as the method used in SPM (Worsley & Friston, 1994; Flandin & Friston, 2017), the accuracy of these formulae depends on the spatial smoothness of the noise in the underlying data. An alternative to using an asymptotic formula is a simulation approach, as commonly used in AFNI (Cox, 1996). In this approach, datasets made up of computer-generated “noise” are used to determine the rate of cluster detection. The cluster-simulation approach avoids the pitfalls of the formula-based approach, at the cost of significant brute force computation. However, performance of early iterations of the cluster-simulation approach were dependent on S. Recently, (Cox et al., 2017b), AFNI’s parametric cluster-simulation approach was improved by modeling the noise spatial autocorrelation function (ACF) with a non-Gaussian shape—termed a “mixed model” ACF, as it combined Gaussian and mono-exponential components—and were able to improve the FPR performance markedly. However, the FPR performance was still dependent on the choice of S, and only for relatively stringent per voxel *p*-thresholds (≤0.002) did the modeling produce anything like the desired global false positive rate. Further development of a separate nonparametric analysis, which omitted any explicit model for the ACF, was able to achieve robust FPR control for a wider range of S, extending to voxelwise *p*≤0.01 by combining a pure *t*-test resampling and cluster-simulation method. Unfortunately, the penalty for this latter approach to detection is that the resulting C thresholds can be very large for studies with relatively few subjects. In addition, the false positives are not uniformly distributed in the brain volume even for these kinds of approaches (Eklund et al., 2016), suggesting that biases still remain, in large part due to regional differences in smoothness across the brain.

ETAC works by implementing a collection of related individual component tests—called *sub-tests* from here onwards—across a range of parameter values, such as smoothing radius and voxelwise *p*-threshold, for dual (voxelwise S, then cluster-wise C) thresholding, and then merging the surviving clusters from the collection of sub-tests. Each sub-test is a “standard” type of test carried out in FMRI group analyses, as exemplified in (Eklund et al., 2016). The ETAC method maintains *equity* (or balance) among this collection of sub-tests, and therefore across otherwise multiple arbitrarily chosen parameters, by constraining each individual sub-test’s cluster-threshold to produce the same FPR as every other sub-test. The final FPR is controlled by adjusting all subtests’ equitably constrained cluster-threshold parameters simultaneously, in order to reach the desired target FPR in the simulations and final results.

ETAC’s use of multiple sub-tests reduces the number of arbitrary choices the FMRI data analyst must make. Effectively, multiple *p*-thresholds and multiple levels of data blurring are all included, and the data analyst only needs to choose ranges of these parameters, rather than a specific *p*-threshold and a specific spatial blur—each combination comprises one sub-test. As a result, ETAC has the potential to detect both small, intense clusters (found using small *p*-value thresholds and small blurring) and large, weak clusters (found using large p-value thresholds and perhaps more blurring) within a single execution. This choice of sub-tests is intended to balance cluster detection across spatial scale (cluster-size) and cluster intensity. As will be demonstrated, ETAC still gives reasonably accurate control of false positives and provides similar statistical power compared to the use of a fixed cluster-size threshold which also controls global FPR properly. Thus, the benefits of equity do not appear to require a tradeoff in FPR or power; the main tradeoff seems to be in having greater computational demands (because of the necessity of multiple sub-tests); but as discussed later, even these are not extreme for typical FMRI studies.

In the next section, the concepts underlying the ETAC approach are outlined. In the Appendix, I provide a detailed summary of how ETAC is implemented in AFNI. To validate the performance of ETAC, I examined both the FPR and the power to detect real activations. To examine the rates of ETAC allowing false results to survive correction, I used simulations *à la* (Eklund et al., 2016), with resting state FMRI as null data for task-based analyses. To examine the ability of ETAC to detect real results (i.e., investigating its utility), I analyzed a publicly available collection of task FMRI datasets and compared the findings to existing methods of multiple comparisons correction which also control global FPR.

## METHODS

### Conceptual Description of Approach

Figure 1 shows a schema for judging all possible voxel clusters in a brain mask (i.e., globally). A point (*S,C*) in this configuration space represents a cluster of *C* neighboring voxels, each of which is individually more statistically significant than *S* (standing in for *t*, *z*, −log(*p*), or similar statistic). The dual thresholding approach for finding significant clusters is also shown: first, a voxel threshold is chosen (a value along the abscissa, e.g., *S*\*), which delimits part of the configuration space; one then names a desired global—within brain mask—FPR *ϖ* (cursive Greek “pi”) and determines an appropriate cluster-size *C*\*, which further delimits the configuration space to the dark shaded region. Thus, *C*\* depends on both S and *ϖ*. For a given global FPR, the cluster-size threshold *C*\*=*C*(*S*\*, *ϖ*) is a decreasing function of per-voxel threshold statistical significance *S*\*, as shown. There is a tradeoff implicit in the dual thresholding approach: a more stringent per-voxel threshold (moving the shaded region rightwards) allows for detection of smaller clusters (the shaded region extends further downwards). The function *C*(*S,ϖ*) is determined by simulation in AFNI and by an asymptotic formula in SPM.

**Figure 1.**
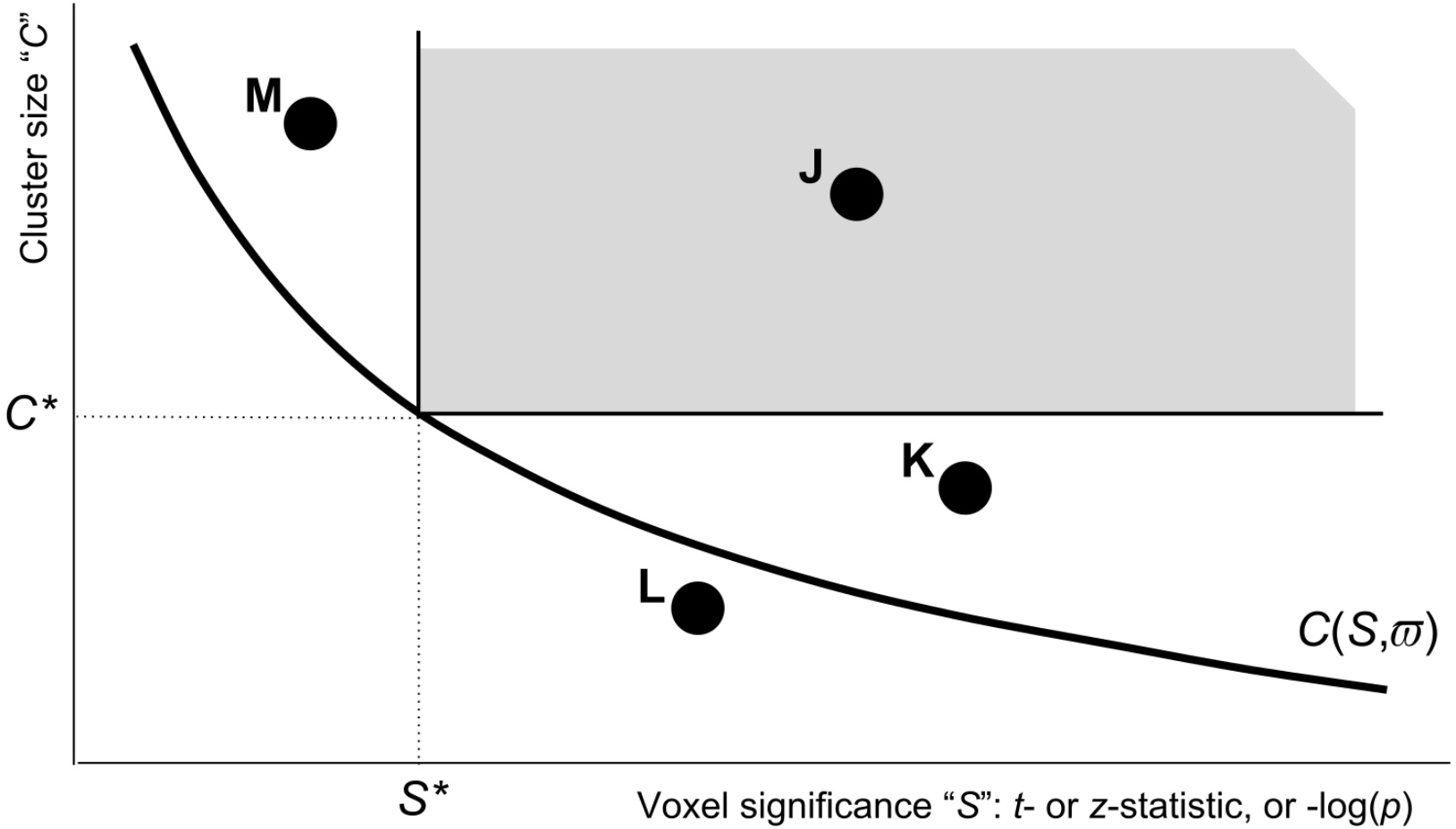
Graphical view of the dual thresholding tradeoff. All voxels that meet a voxelwise threshold and a cluster-size (contiguity) threshold are kept—the gray region indicates the combinations that pass a particular instance of this procedure: each voxel’s statistic must pass a first significance threshold S*, and the number of contiguous voxels (in a single cluster) must be above a second threshold C*. The thick black curve C(S,*ϖ*) indicates the cluster-size threshold that gives a fixed global FPR *ϖ* (e.g., 5%) as a function of the per-voxel threshold S; *ϖ* is the probability that a noise-only cluster falls into the gray region. See the main text for discussion of named clusters J-M.

In Figure 1, the J cluster passes the thresholding test, by containing enough voxels for the given voxelwise threshold. The K cluster is made up of voxels that easily pass the threshold test, but it does not have enough voxels to survive in the present scenario; however, if the per-voxel threshold test were more stringent (*S*\* moved rightward), then the K cluster could pass the dual threshold test. For the right choice of per-voxel threshold, both clusters J and K would survive. The L cluster has voxels that pass the per-voxel threshold, but not enough of them and there is no per-voxel threshold that will permit L to survive at the desired FPR. Conversely, the M cluster has a lot of voxels above a slightly smaller per-voxel threshold. Again, one could make M survive by playing with the per-voxel significance threshold, as M lies above the *C*(*S*, *ϖ*) tradeoff curve. For a given nominal FPR *ϖ*, the choice of voxelwise thresholding statistic *S*\* is essentially the primary quantity for determining which clusters are accepted or rejected, because the cluster-size threshold *C\** is a function of *S*\* parametrized by the *ϖ* value (e.g., 5%).

Figure 1 and the reasoning above lead to the obvious question: since researchers are interested in obtaining clusters based on their dual properties of voxelwise significance and size, how should they choose the single voxelwise threshold that determines the result? Previous work (Eklund et al., 2016; Cox et al., 2017b) has demonstrated the bounds of a reasonable range of choices (at some larger *p*-thresholds the ability to control FPR decreases; in the present language, for some large *p*-values (small *S*-values), such as *p*≥0.10, an appropriate *C*(*S*,*ϖ*) may not exist, or may be so large as to be useless). But even within a reasonable range of p-values, the choice of threshold might affect the outcome dramatically. As illustrated in Figure 1, one could shift this voxelwise threshold along the horizontal axis and obtain the following combinations of statistically significant clusters: only M; both J and M; only J; both J and K; or only K. When the results can depend strongly on the choice of an arbitrary parameter, any value of which might considered reasonable within a large interval, the researcher is in a bad situation—as are the readers of any resulting scientific paper.

Figure 1 also leads to the idea of finessing the tradeoff between cluster-size and per-voxel thresholds; that is, simply accept every voxel configuration which lies above the solid black curve *C*(*S*, *ϖ*). From the viewpoint of the final FPR of the dual threshold method, any place along that curve is as acceptable as any other. The principle of equity asks, “Why discriminate?”; however, a practicable algorithm must use a discrete set of *p*-thresholds combined with their corresponding cluster-size thresholds. Finally, the algorithm must be constrained by the need to achieve the desired global FPR in the final results.

Figure 2 illustrates the use of four different per-voxel thresholds within a single cluster test. In this example scenario, using the equitable thresholding method, clusters J, K, and M survive. When using the multiple voxelwise thresholds (i.e., the proposed multi-*S* or multi-*p* case), it is important to note that one cannot use the original *C*(*S*,*ϖ*) tradeoff curve (from the mono-*S* case) to calculate the cluster threshold for each sub-test; this would lead to a final FPR that is larger than the desired *ϖ*, since more potential false positive voxel configurations are allowed with the use of multiple *S*-thresholds than with a single *S*-threshold. It is thus necessary to find an *ϖ*_✭_ < *ϖ*_G_ (subscript “G” for final goal) such that using the *C*(*S*, *ϖ*_✭_) for the individual sub-tests will achieve the desired *ϖ*_G_ in the final combined simulations. That *ϖ*_✭_ value then defines the multi-*p*-threshold clustering algorithm when applied to the original voxelwise statistical tests. ETAC does *not* make the Bonferroni correction *ϖ*_✭_=*ϖ*_G_/4 among these four sub-tests. This correction would be very conservative and negatively impact the power of the method, as indicated by the strong overlap among the individual mono-*p*-threshold regions in Figure 2. The Appendix contains a detailed description of how *ϖ*_✭_ is adjusted (via simulation) to make the final FPR equal to *ϖ*_G_ in the Appendix.

**Figure 2.**
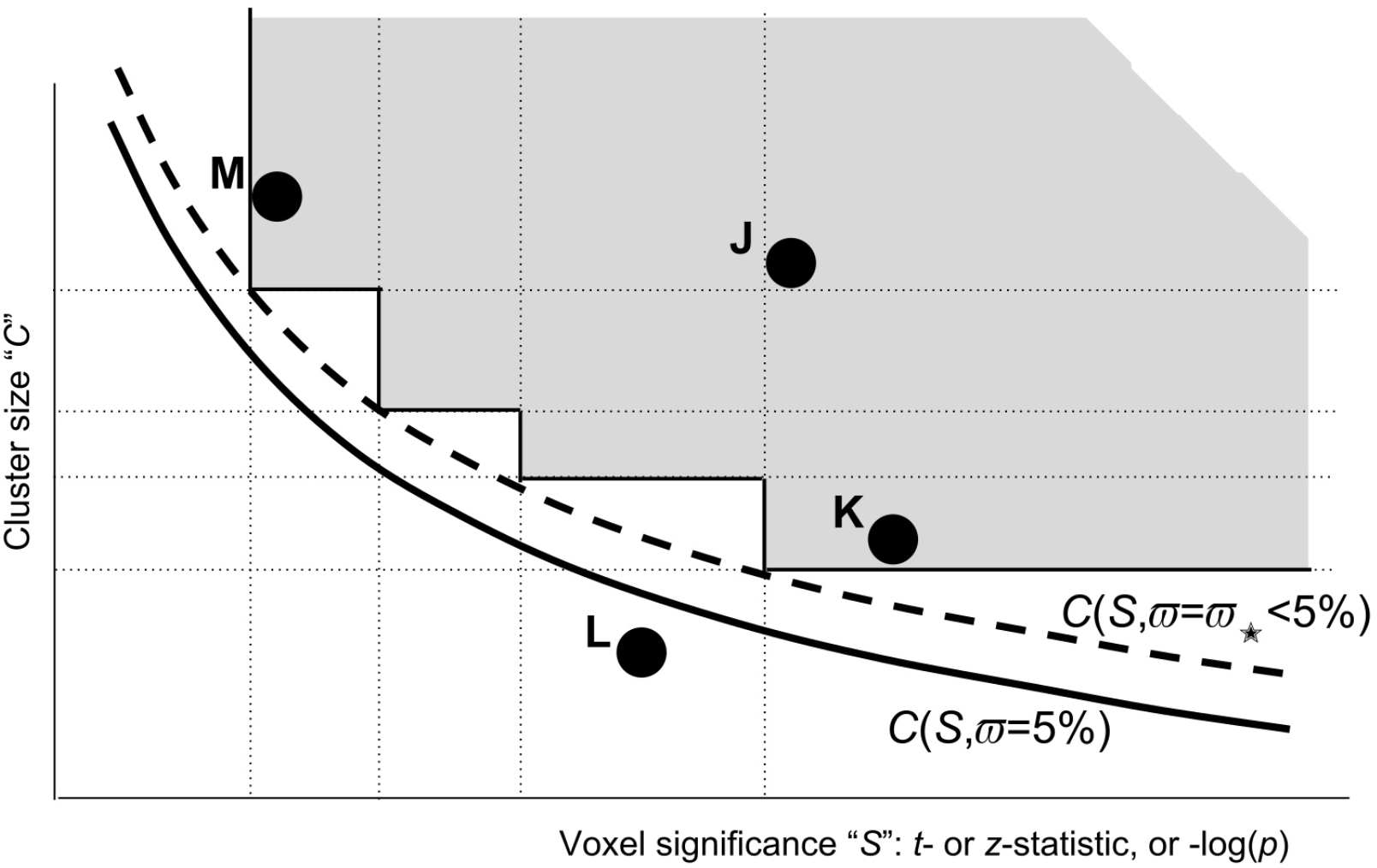
Thresholding as in Fig 1, but using four different per-voxel p-thresholds at once (vertical dotted lines), each using its own cluster-size threshold (horizontal dotted lines). The FPR for the gray region (the union of four mono-p-threshold rectangles), if constructed above the solid black curve C(S, *ϖ*), would be larger than the desired *ϖ* (FPR for the gray rectangle shown in Fig. 1), since this union region would include more possible voxel configurations. To compensate for that and maintain a nominal (5%) FPR, the cluster-size threshold curve from Fig. 1 (solid curve) must be raised to a higher level by picking a curve C(S, *ϖ*_✭_ with an *ϖ*_✭_ < 5% (dashed). Which *ϖ*_✭_ should be chosen is determined by simulating this multi-p-thresholding process—just as the curve C(S,*ϖ*) for the original dual threshold method is determined (in AFNI) by simulating the mono-p-threshold procedure. (cf. Fig 6.)

A key advantage of this approach over the simpler mono-*S*-thresholding method is that it allows large clusters of low effect size (smaller *S*, larger *p*) to be detected along with small clusters of high effect size (larger *S*, smaller *p*) within a single analysis. In the more common mono-*p*-threshold approach, the user (or software) has to decide which type of cluster to favor. It would be difficult for a user to know *a priori* what single *p*-value would be appropriate or preferable for a study. This arbitrariness could all too easily lead to the user adjusting the per-voxel *p*-threshold to get more desirable results. Additionally, a user may be interested in both small and large clusters (e.g., responses in amygdala and posterior cingulate cortex) within a single experiment, an outcome which might be arbitrarily excluded by the constraint of being allowed only a single voxelwise threshold. In the ETAC method, only a plausible range of *p*-thresholds needs to be chosen, and these issues are effectively removed. In summary: where results initially depended on just a single parameter chosen from within a range of reasonable values, the ETAC method allows multiple parameter values to be chosen, tested, and have their results combined into a single result, while simultaneously balancing the importance of each subtest and maintaining control of the final FPR.

The idea of combining multiple dual threshold sub-tests for detection (in Fig 2, each subtest is specified by its voxelwise *S*- or *p*-threshold) can be directly abstracted and generalized to other parameter dependencies, such as the scale of the spatial smoothing applied to the data (parameterized by the full-width at half max, FWHM, of a Gaussian shape). Similar to S, typically only a single blur size is chosen for a given FMRI study, but commonly implemented values within the literature range over 4-10 mm. No matter what the sub-test is, its individual FPR will be controlled by a cluster-thresholding parameter. The ETAC approach is to require each sub-test to have the same individual FPR *ϖ*_✭_ and to accept a voxel if it passes *any* of the individual sub-tests (set union); then, *ϖ*_✭_ is adjusted to achieve the goal of FPR=*ϖ*_G_ for the final set of accepted voxels. In this way, equity/balance across the parameter ranges of the sub-tests (in Fig. 2, defined by *S* threshold values) is maintained while the overall FPR is still achieved.

Using multiple cases of blurring to enable FMRI detections across spatial scales is certainly not new (Worsley et al., 2001). The ETAC framework maintains equity across blur scales in the same way as described above: the clustering thresholds at the different scales of blurring are balanced by making each sub-test (i.e., combination of *p*-threshold and blurring FWHM) have the same individual FPR *ϖ*_✭_. The effect of this multi-blur analysis is to remove the arbitrary choice of blurring scale from the data analyst. For this approach to work, the first-level (individual subject) analyses should be done *without* spatial blurring. Then, the blurring of the first-level results (individual subject betas; in AFNI, beta estimates percent signal change from voxel baseline) should be carried out by the ETAC software.

### Computational Approach

Building on the methods of (Cox et al., 2017b), ETAC builds a large collection of noise-only *t*-statistic volumes using randomization/permutation of the original input datasets. These volumes are then thresholded, clusters are formed, and various scores for each cluster are computed. ETAC false positive rates for this large collection of simulated volumes and clusters are computed (see Appendix for algorithmic details) and adjusted until the desired global FPR is met. Creating the collection of noise-only volumes by repeatedly *t*-testing thousands of randomized cases is the lengthiest part of the computation.

Cluster size is a very widely used score or figure of merit (FOM) for assessing significance in neuroimaging. The size of a cluster (voxel count) is the sum of the number 1 across all voxels within the cluster; that is, across *N* voxels, **Σ** 1. From this formulation, it is simple to generalize the cluster significance FOM to other sums across cluster voxels, sometimes referred to as “cluster mass” (Zhang et al., 2008); for example, incorporating the statistic associated with each *i*-th voxel as a weight, such as 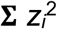, where *z_i_* denotes the 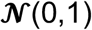 normal deviate that matches the voxel-level test statistic (e.g., *t* or *p*) in tail probability. The current implementation of ETAC in AFNI allows for the cluster-thresholding FOM to be computed as **Σ** |*z_i_*|^*h*^ where *h*=0, 1, or 2; the h=0 case corresponds to the unweighted cluster size (i.e., the sum of 1s), while *h* > 0 provides for statistical significance weighting, which allows for a somewhat increased possibility of finding smaller but more intense clusters. (After some experimentation, *h*=2 was chosen as the default.)

## TESTS and RESULTS

### Tests 1: FPR control

The first test of the proposed ETAC approach uses a variation on the methodology in (Eklund et al., 2016). From the FCON1000 collection of resting-state FMRI datasets (Biswal et al., 2010), the 198 datasets in the Beijing sub-collection were processed with AFNI pipelines specified using afni_proc.py to produce the first-level (individual subject) activation maps. In this process, datasets were resampled to 2 mm^3^ voxels to conform with (Eklund et al., 2016); this refinement is not needed for any other reason (the original voxel dimension is about 3 mm). The Beijing sub-collection was chosen for this test to test ETAC in a similar setting as used in (Eklund et al., 2016)—the paper that launched this current effort. Five sets of pseudorandom timings were used to produce 198×5=990 maps of fit coefficients (betas), with no spatial blurring used at this analysis level. These 990 maps were each computed for three different pseudo-task stimulus durations: 1 s, 10 s, and 30 s. The BOLD hemodynamic response function (HRF) regressor models for all 5 pseudo-task timings in each duration were nearly orthogonal—stimulus duration 30 s had the most highly correlated HRF models, with mean correlation 0.09 and standard deviation 0.21—so that the 5 estimated response magnitudes (betas) in each subject were approximately uncorrelated. This use of multiple randomized timings slightly distinguishes the present testing methodology from that in (Eklund et al., 2016).

At the second (group) level, two-sample *t*-tests were carried out, with 20 randomly chosen subjects (of the 198) assigned to each sample. For each subject, 1 of the 5 results from the variable task timings was selected randomly in each simulation. A total of 1,000 realizations of this procedure was analyzed with ETAC, for each of the 3 stimulus durations. For each stimulus class (1s, 10s, 30s), three results were calculated: the FPRs for 1-sided (positive or negative) or 2-sided *t*-thresholding; thus, there are 3×3=9 results for the two-sample *t*-tests for each goal FPR. A very similar set of simulations was run separately using one-sample *t*-tests with 40 subjects per simulation. In all cases, a panoply of nominal (goal) FPRs was run, with *ϖ*_G_=1%, 2%, …, 9%.

Within the ETAC procedure, 3 blur cases were used in defining the sub-tests: 4 mm, 7 mm, and 10 mm (FWHM). Ten voxelwise *p*-thresholds were used, 0.010, 0.09, 0.08, …, 0.001 (the defaults); thus, 30 sub-tests were used in each of these simulations. The NN cluster-defining level was left at the default 2 (i.e., neighbors of a voxel are those sharing either face or edge; cf. Appendix), and the default FOM with *h*=2 (i.e., statistically weighted) was used.

The empirical FPRs from 1-sample and 2-sample *t*-test simulations are presented in Figure 3 for the ETAC method, along with the 95% (binomial) confidence intervals that would be expected from 1,000 *independent* simulations if the algorithm exactly achieved the nominal FPR. For the two-sample *t*-tests (with 40 subjects total), the results are tightly clustered, slightly below the nominal FPR and typically within the confidence interval. For the one-sample tests with 40 subjects, the results are more variable. This effect is due to the lack of independence among the resampled dataset collections, as each collection of 40 is taken from the same 198 source datasets; see also (Eklund et al., 2016; supplementary material). The same effect (broadened variability of achieved FPR) exists in the two-sample tests, but is much smaller.

**Figure 3.**
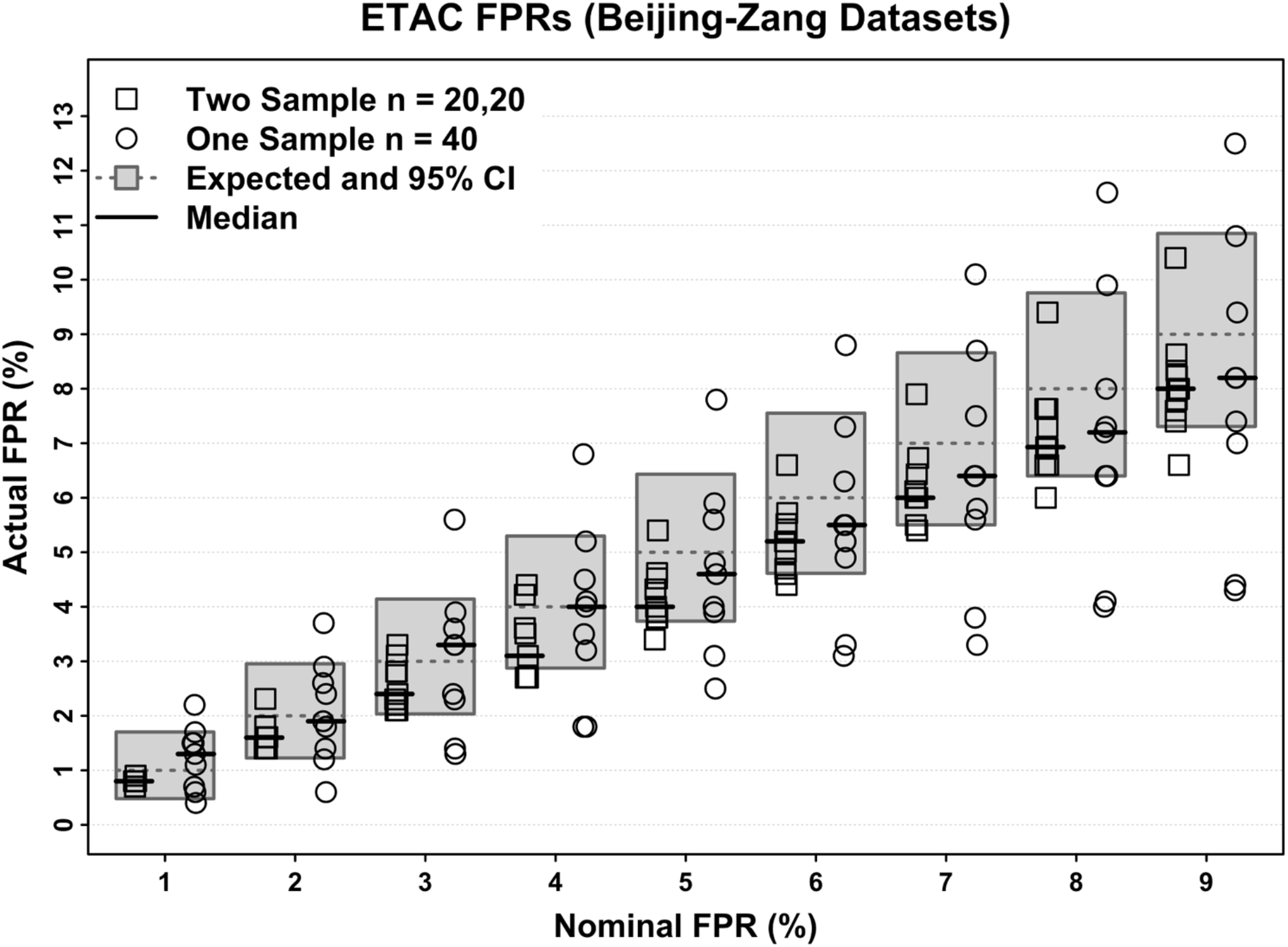
ETAC false positive rates from simulations of two-sample *t*-tests (20 subjects per sample) and one-sample t-tests (40 subjects in the sample). The nominal (desired) FPR *ϖ*_G_ is shown along the horizontal axis, and is also shown as a dashed line within the gray boxes, which represent the 95% confidence intervals for the FPR (from a binomial distribution, assuming the nominal FPR as the binomial parameter and assuming all 1000 replicates are independent). Thus, symbols that fall inside their gray box are within the 95% CI of what would be expected if the nominal FPR was the actual ETAC FPR. Medians for the 9 values in each partition are shown with solid black lines. The two-sample ETAC results are fairly close to the nominal FPR and tightly clustered, but the one-sample results have a great deal more variability; see the main text for an explanation of this increased dispersion.

The two-sample FPR results appear to be slightly biased below their nominal goals. The reasons for this are unknown to the author.

### Tests 2: FPR spatial variability

To examine the spatial distributions of false positives found by ETAC, 10,000 simulated two-sample cases were run using the resting-state FMRI null test methodology described earlier, using the beta maps from the 10s duration pseudo-stimulus timings, with the same 3 blurring parameters and 10 *p*-thresholds. Figure 4 shows the spatial distribution of false positives for various sagittal slices, for ETAC and for mono-*p*-thresholds applied to the individual blur cases (each of the latter were thresholded at *p*=0.005). While, as shown above, the ETAC approach is successful at controlling the global FPR, the cluster results are not uniform across the brain; this result is similar to that previously observed in various cluster-thresholding methods (Eklund et al., 2016; cf. Supplementary Fig 18 et seq.). My goal of producing a markedly more uniform distribution of false positives was not achieved.

**Figure 4.**
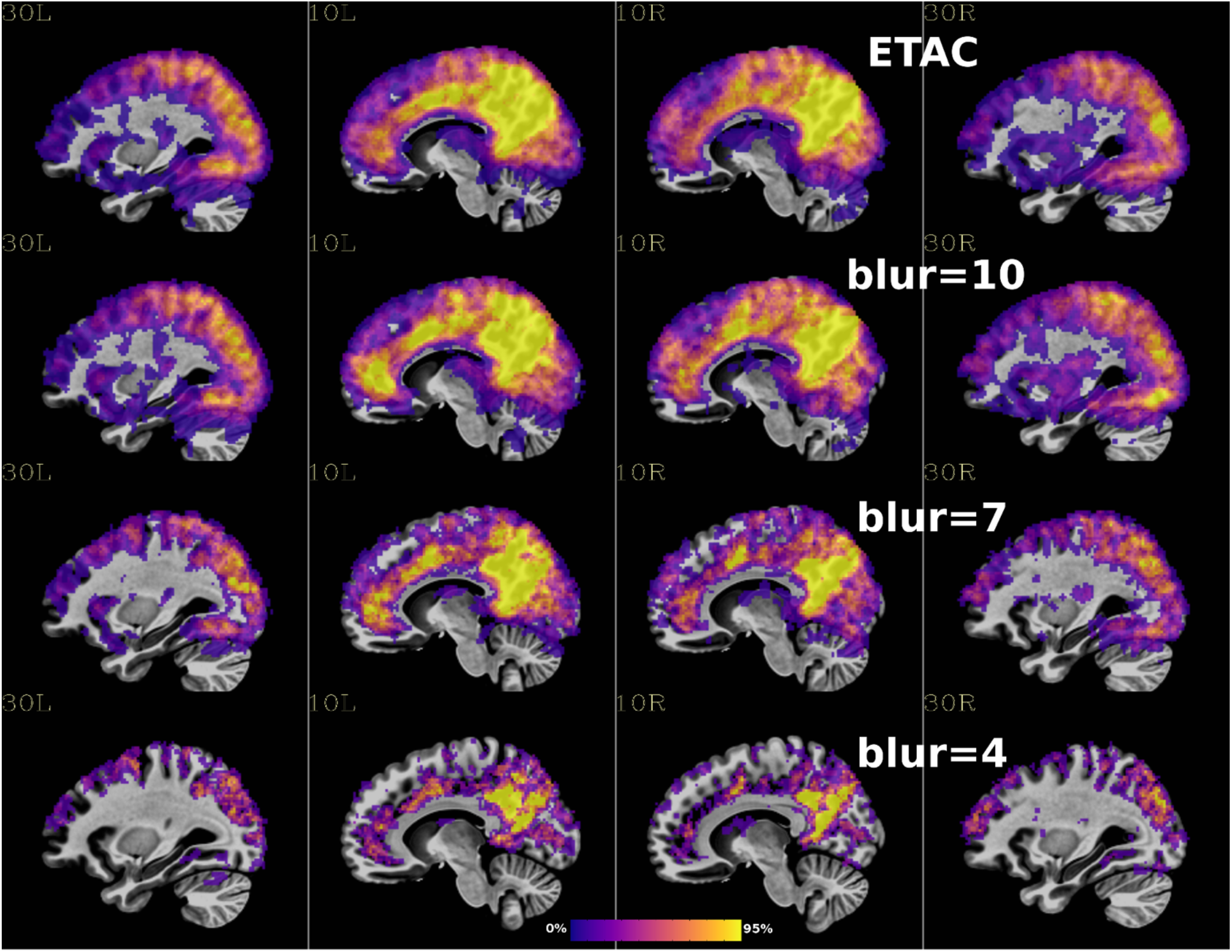
Spatial distribution of false positives resulting from 10,000 null simulations. ETAC was run with *ϖ*_G_=5%, 10 p-thresholds (0.001, 0.002,… 0.010), 3 blurs (4mm, 7mm, 10mm); ETAC’s results are mapped in the upper panel. The lower 3 panels show the spatial distributions for the single p=0.005 threshold method for each blur case. Since the range of false positive counts varies strongly with blur case, the color range for each panel is different and is set to the 95% point on the cumulative distribution of the nonzero FPR voxel counts in each case.

### Tests 3: Reliability

To examine the ability of ETAC to find true positives, a collection of task datasets was downloaded from OpenFMRI.org. The UCLA Consortium for Neuropsychiatric Phenomics LA5c Study (Poldrack et al., 2016; OpenFMRI collection ds000030) was chosen, in particular the pattern matching and encoding “pamenc” task, using 78 control subjects (four control subjects’ data had to be discarded due to excessive head motion). This collection of data was selected since it is a relatively large study, and the pamenc task robustly engages large segments of the brain involving vision, spatial perception, and language. Two-sample *t*-tests were carried out with 20 subjects per sample, randomly selected from the 78. Sample A was drawn from the pamenc task results and sample B from the control task results (separate random subject lists for each sample of 20). This analysis was carried out with 1000 replicates with ETAC, using *h*=0 (cluster-size thresholding), with *ϖ*_G_ =5%, and also using several mono-*p*-thresholds. In this analysis, only one blur level with FWHM=7 was used, to allow direct comparison of results between ETAC (with 10 *p*-thresholds) and mono-*p*-threshold results.

Figure 5 shows the map of the *differences* in the detection rate between ETAC and mono-*p*-thresholding at *p*=0.005 and *p*=0.010. (Figure 5 is *not* an activation map, but instead shows the places where the two methods differed in likelihood of finding results). Comparing ETAC to mono-*p*=0.005 (upper panel), it is clear that ETAC’s results cover significantly more volume. Comparing ETAC to mono-*p*=0.010 (lower panel), there are fewer differences and they are in both directions; ETAC finds some more results in places, and few in others.

**Figure 5.**
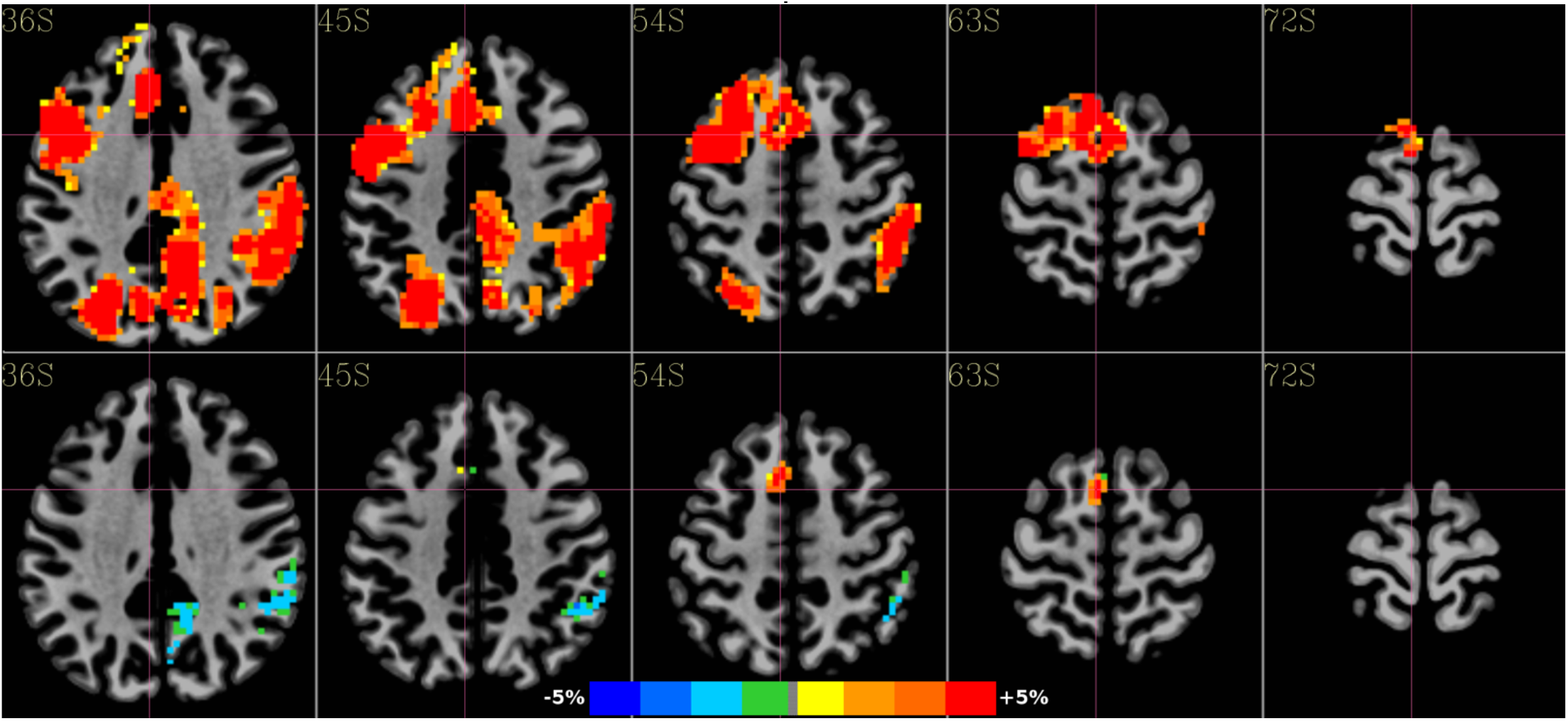
Maps of the *differences* in the detection rate between ETAC and mono-*p*-thresholding at *p*=0.005 (upper) and *p*=0.010 (lower), calculated from running 1,000 simulations with the UCLA phonemics study, pamenc-vs-control tasks *t*-test. Color range is set to 5%; for example, red means that ETAC detected a result at least 5% more often than the mono-*p*-threshold method, and dark blur indicated that the mono-*p*-threshold method detected a result at least 5% more often than the ETAC method. (Much more of the brain is truly active than shown here; in many places, the detection probabilities of the two methods were nearly identical.)

Figure 6 shows a graph of the cluster-size thresholds involved in computing Figure 5 (for *ϖ*_G_ =5%), and is a real-life instantiation of the conceptual Figure 2. The lower curve (triangles) shows the cluster-size thresholds needed for mono-*p*-thresholding. The upper curve (circles) shows the cluster-size thresholds used with ETAC’s 10 *p*-thresholds. The boxes-only plot shows the cluster-size thresholds resulting when ETAC is re-rerun specifiying 91 *p*-thresholds from 0.001 to 0.010 (Δ*p*=0.0001). The 10 *p*-thresholds cluster-sizes faithfully follow the 91 *p*-thresholds plot, and are raised above the mono-*p*-thresholds cluster sizes. At *p*=0.010, the ETAC cluster-size threshold is 1383 voxels while the mono-*p*-threshold is 1053 voxels. This greater stringency at *p*=0.010 is why Figure 5 shows places where ETAC is somwhat less likely to find clusters than mono-*p*=0.010. The tradeoff is that ETAC can find clusters where the mono-*p*-threshold method cannot.

**Figure 6.**
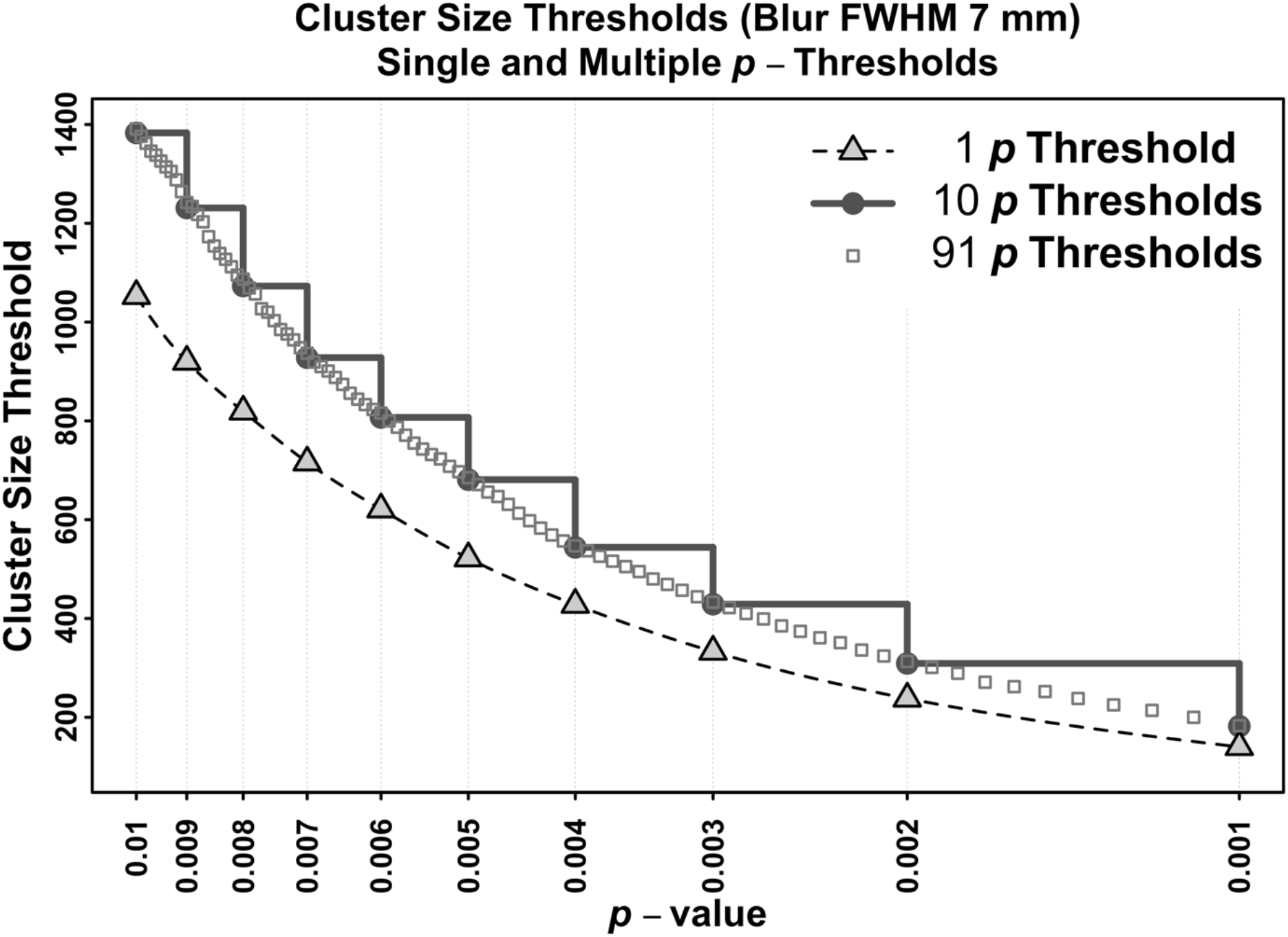
Cluster-size thresholds involved in computing Fig 5 (for *ϖ*_✭_=5%). The lower curve (triangles) shows the cluster-size thresholds needed for mono-*p*-thresholding. The upper curve (circles) shows the cluster-size thresholds used with ETAC’s default 10 *p*-thresholds. The boxes-only plot shows the cluster-size thresholds resulting when ETAC uses 91 *p*-thresholds from 0.001 to 0.010 (Δ*p*=0.0001). (cf. Fig 2.)

## DISCUSSION

The ETAC method has been shown to be effective at controlling the false positive rate (FPR) for cluster-based detection in task FMRI. This performance is achieved while also reducing the influence of the arbitrary selection of parameters affecting the cluster results, the primary motivation for the method’s inception. Importantly, ETAC does not lose much true detection power relative to simpler non-parametric cluster-thresholding methods, and can have more power depending on the study and on the simpler method chosen for comparison.

In the testing framework used in (Eklund et al., 2016) and in (Cox et al., 2017b), non-parametric clustering methods were shown to control FPR better than parametric methods that are based on a model for the spatial correlation function of the FMRI noise. These non-parametric methods, though, are more stringent than the parametric methods, having reduced power by comparison, and will lose some results that parametric methods will detect. In other words, the usual statistical tradeoff is still present. ETAC’s use of multiple sub-methods can recover a little of the lost statistical power, as evidenced in Fig 5, but it is still more strict than the parametric clustering methods in widespread use since the cluster-size/FOM thresholds are usually larger than those the parametric models yield. A noteworthy advantage of ETAC is that it removes two arbitrary choices from the group analysis: the voxelwise *p*-threshold, and the amount of spatial blurring to apply. For example, there are no strong *a priori* reasons to favor *p*=0.001 over *p*=0.01 (provided FPR is properly controlled), or 8 mm (FWHM) blurring over 4 mm blurring.

The computational burden of the full ETAC method is not especially onerous, since it is run on all of the data in a study to produce a group map. For instance, an ETAC run with 40 subjects at 2 mm^3^ resolution, with a single goal FPR *ϖ*_G_, ran for 42 minutes on a 16 core Intel/Linux node, and needed 17 gigabytes of memory to hold all the intermediate simulations and cluster tables. Most of this time and memory is needed for the repeated *t*-tests of the randomized/permuted input datasets. Expanding the *ϖ*_G_ list to the entire gamut of 1%, 2%, …, 9% increased the run time to 51 minutes.

While the computational cost of performing the ETAC analysis is not prohibitive, it could make it difficult to extend the ETAC method to more complex voxelwise statistical mapping techniques, such as Linear Mixed Effects analyses (Chen et al., 2013). Such analysis methods are themselves computationally intensive, unlike simple voxelwise *t*-tests or linear regression. Thus, iterating and randomizing such tests thousands of times is at present impracticable.

## CONCLUSION

I have described the ETAC (Equitable Thresholding and Clustering) methodology and shown comparisons of it with an existing standard cluster-threshold technique that also controls false positive rates reasonably well. ETAC’s primary benefit is to reduce the dependence of arbitrary parameter choices on final cluster results, as well as to allow for the discovery of a greater range of cluster types (large with relatively low voxelwise significance or small with relatively high voxelwise significance) in a single analysis and with essentially equal footing.

## Acknowledgements

The research and writing of the paper were supported by the NIMH Intramural Research Programs (ZICMH002888) of the NIH/DHHS/USA. This work would have been impossible without the outstanding computational resources of the NIH HPC Biowulf cluster (http://hpc.nih.gov). Data for the analyses included the FCON-1000 collection (https://www.nitrc.org/projects/fcon_1000/) and OpenFMRI collection ds000030 from (https://openfmri.org/dataset/ds000030/). Thanks go to Paul Taylor for many suggestions that improved the manuscript immeasurably. Thanks go to the referees for very useful critiques (p<0.01). The ideas for equitable application of multiple sub-tests came to RWC while hiking in Grand Canyon National Park; thanks go to the US National Park Service for maintaining this inspirational public treasure.

## ACRONYMS

ACF: Autocorrelation Function
AFNI: Analysis of Functional NeuroImages
BOLD: Blood Oxygenation Level Dependent
EPI: Echo Planar Imaging (or Image)
ETAC: Equitable Thresholding and Clustering
FMRI: Functional Magnetic Resonance Imaging (or Image)
FOM: Figure of Merit
FPR: False Positive Rate
FWHM: Full Width at Half Maximum
HRF: Hemodynamic Response Function
SPM: Statistical Parametric Mapping

## Appendix Implementation Details

ETAC is implemented inside the AFNI group analysis program 3dttest++, for convenient use by the data analyst. All operations take place inside a user-selected mask of voxels (e.g., a brain mask or a gray-matter mask). Although the generic concept of balancing detection across multiple methods is relatively straightforward, the actual implementation is mildly complex, as many details need to be accounted for along the way. An outline of the procedure is:

A. Generate *N* (default: 10,000) noise-only random fields (volumes) of *z*-statistics for cluster analysis; the following steps are used to create each random field generated:

a. The residuals in the voxelwise GLM (*t*-tests, possibly with subject-level covariates) are sign-randomized; if two subject samples are used, the subjects’ datasets are also randomly permuted between groups (unless covariates are present).
b. The GLM is repeated using the randomized/permuted sample, saving the resulting voxelwise *z* statistics (converted from *t*).
c. The program requires at least 17 input datasets to allow a reasonable number of randomizations.
d. These *N* realizations can also be used for the nonparametric cluster-size threshold analysis described in (Cox et al., 2017b), which will produce a table of cluster-size thresholds vs. *p*-thresholds; essentially, the *C*(*S*,*ϖ*) curve from Fig. 1 (generated for a range of *p* and *ϖ* values).
B. Apply each of the sub-tests (i.e., tensor product of *p*-thresholds, blur cases, and *h* values):

a. For each sub-test, form all clusters in all *N* random fields.

i. The default *p*-thresholds are 10 values distributed linearly between 0.01 and 0.001 (0.010, 0.009, …, 0.001). The data analyst can choose a different set of *p*-thresholds. In practice, there is very little difference between using 10 and using 91 equally spaced *p*-thresholds over the same interval (cf. Fig 6).
ii. There is no default set of blur cases; rather, the default is not to apply additional blurring, since the ETAC group analysis software doesn’t “know” how much blurring was applied to its input data during preprocessing. If multiple blur cases are specified, each blur case has the entire set of *p*-thresholds applied.
b. Save a table of all clusters found for each sub-test. Each cluster data structure includes a list of its voxels, and its computed FOMs (for *h*=0, 1, 2). The default FOM is 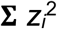 (e.g., *h*=2).
C. <u>Combining sub-tests</u>:

a. For each sub-test and for each cluster-FOM type computed, create a sorted array (largest first) of the cluster-FOM values produced in the simulations. These arrays represent the reversed cumulative distribution functions of the cluster-FOM statistics under the global null hypothesis (no significant group results anywhere).
b. Pick a fraction *τ* (initial value (4+*ϖ*_G_)·0.0006), and set the cluster-FOM threshold for each sub-test to be that sub-test’s FOM array’s (*τN*)’th largest entry; that is, a larger *τ* selects a smaller threshold (moving down in the sorted array). For example, when *N*=10,000 and *ϖ*_G_=5, the initial cluster-threshold for each sub-test is the 54^th^ entry in the corresponding array. (If *τN* is not an integer, the cluster-threshold is interpolated from the sorted FOM array.) *This uniform selection in the FOM lists is how equity is applied across sub-methods, since each FOM list is from one sub-method*.
c. Apply the resulting cluster FOM threshold values to produce the significant voxels map for each of the *N* random fields, merging the results from each sub-test, and count the fraction *φ* of the *N* random fields that have *any* surviving voxels; *φ* is the estimate of the global FPR.
d. Adjust the fraction *τ* up (if *φ*<*ϖ*_G_) or down (if *φ*>*ϖ*_G_), until is *φ* approximately at the target *ϖ*_G_ (which may be any value from 1% to 9%; the default, of course, is 5%).

i. If any two previous *φ* results bracket *ϖ*_G_, then inverse linear interpolation in *φ*(*τ*) is used to adjust *τ* for the next trial.
ii. Otherwise, *τ* is just scaled linearly by *ϖ*_G_/ *φ* for the next trial.
iii. In usage thus far, usually only a few (2-3) iterations are necessary for convergence.
iv. The user has the option to compute threshold maps for a collection of FPR goals *ϖ*_G_=1%, 2%, …, 9%, in order to allow perusal of the results at various levels of statistical stringency.
e. Apply the final multi-method cluster-FOM threshold maps to the actual *t*-statistics resulting from the original GLM analysis.

i. If multiple blur cases are used, each blur case’s multi-*p*-threshold maps are applied to the GLM tests on blurred copies of the original individual-subject datasets, and the set of resulting detection maps (one from each blur case) are merged to make the final detection map.
ii. The outputs are a binary map (NIFTI dataset), indicating which voxels survived the process. This dataset can be used to mask the original GLM results to produce a final activation map. A second dataset is also produced, indicating which of the sub-tests were passed for each voxel. The main use for this dataset is for analyzing the ETAC process itself, to determine which sub-tests might have contributed unique results. A table of the cluster-FOM thresholds is also saved, of course (after all, this table was one of the principal goals of the extensive number crunching).

The software is written in the compiled language C to be able to run in an acceptable time frame for group analyses, and is parallelized to use multiple CPU cores. In typical cases, the majority (70+%) of the computational time is spent carrying out the 10,000 repeated *t*-tests to produce the simulated noise volumes.

Cluster-contiguity can be defined in several ways. In AFNI, contiguity is defined by the nearest neighbor (NN) level, which can be 1, 2, or 3:

1. Voxels are defined as contiguous if faces touch (first nearest neighbors);
2. Voxels are defined as contiguous if faces or edges touch (second nearest neighbors);
3. Voxels are defined as contiguous if faces, edges, or corners touch (third nearest neighbors).

The ETAC software in AFNI does *not* allow for balancing across these different clustering possibilities; the NN level remains a parameter the analyst must choose (default value in AFNI’s ETAC is 2). I do not think that providing equity across different clustering methods would be of major consequence.

The application of different blur cases in ETAC is carried out by applying 3D Gaussian blurring to the input datasets, but the kernel is restricted to the voxel mask supplied by the user. This blur-in-mask procedure is carried out by a finite difference pseudo-time stepping method applied to the 3D diffusion equation, with Neumann (reflecting/conserving) boundary conditions at the edge voxels of the mask.

The cluster FOM chosen can be **Σ** |*z_i_*|^*h*^ for *h*=0, 1, and/or 2. The software can balance across any combination of these FOMs; however, the default choice is the single FOM with value *h*=2. In practice, I see little advantage in balancing across multiple cluster FOM formulæ. Other FOM formulæ could be added to the software with relatively little effort.

